# Cannabinoid receptor deficiencies drive immune response dynamics in *Salmonella* infection

**DOI:** 10.1101/2025.03.10.642352

**Authors:** Hailey A. Barker, Saloni Bhimani, Deyaneira Tirado, Leandro Nascimento Lemos, Luiz F.W. Roesch, Mariola J. Ferraro

## Abstract

This study investigated the roles of cannabinoid receptors 1 and 2 (CB1R and CB2R) in regulating host responses to *Salmonella Typhimurium* in C57BL/6 mice. The absence of both receptors significantly impaired host resilience, as evidenced by increased weight loss, deteriorated body condition, and reduced survival following infection. Notably, CB1R deficiency resulted in more pronounced weight loss and heightened susceptibility to bacterial proliferation, as demonstrated by increased *Salmonella* dissemination to organs. In addition, both CB1R and CB2R knockout mice exhibited alterations in immune cell recruitment and cytokine production. CB1R-KO mice displayed increased T cell and macrophage populations, whereas CB2R-KO mice showed a reduction in NK cells, indicating receptor-specific effects on immune cell mobilization. Cytokine profiling of macrophages post-infection revealed that CB1R-KO mice had reduced IL-10 levels, along with increased IL-6 and TGF-β, suggesting a dysregulated polarization state that combines pro-inflammatory and regulatory elements. In contrast, CB2R-KO mice exhibited a profile consistent with a more straightforward pro-inflammatory shift.

Furthermore, microbiota analysis demonstrated that CB2R-KO mice experienced significant gut dysbiosis, including reduced levels of beneficial *Lactobacillus* and *Bifidobacterium* species and an increase in pro-inflammatory *Alistipes* species post-infection. Functional microbiome analysis further indicated declines in key metabolic pathways, such as the *Bifidobacterium* shunt, L-glutamine biosynthesis, and L-lysine biosynthesis, suggesting microbiota-driven immune dysregulation. Together, these findings highlight the distinct, non-redundant roles of CB1R and CB2R in modulating innate immunity, host defense, and microbiota composition during bacterial infections.

**Significance Statement:** Understanding the role of cannabinoid receptors in immune regulation is important for identifying new therapeutic targets for bacterial infections. Our study demonstrates that CB1R and CB2R play distinct, non-redundant roles in host defense against *Salmonella* Typhimurium. The absence of these receptors impairs host resilience, increases bacterial dissemination, and alters immune cell recruitment and cytokine production. Notably, CB1R deficiency leads to enhanced weight loss, increased bacterial spread, and a dysregulated macrophage cytokine profile—characterized by reduced IL-10 and elevated IL-6 and TGF-β—while CB2R deficiency is associated with reduced NK cell numbers and a more pronounced pro-inflammatory cytokine profile. These findings reveal a receptor-specific balance in immune responses, suggesting that cannabinoid signaling modulates infection outcomes. Targeting CB1R and CB2R pathways may offer novel strategies to enhance host immunity and improve treatments for bacterial infections in the future.

## Introduction

Cannabis consumption is increasing in the U.S. for both recreational and medical purposes ^1^, yet its effects on infections remain inadequately understood. Cannabis contains over 113 cannabinoids (CBs), including Delta-9-tetrahydrocannabinol (THC), a partial agonist of the cannabinoid type 1 receptor (CB1R) primarily in the central nervous system, and CB2R, which is abundant in immune cells^2^. These medically relevant cannabinoids include Cannabidiol (CBD), Nabiximol or Dronabinol. CBs interact with the endocannabinoid system (ECS), a neuromodulatory network comprising CB receptors, their ligands, endocannabinoids (eCBs), and the enzymes resp 1onsible for eCB synthesis and breakdown. The ECS functions through the binding of eCBs/CBs to CB1R and CB2R receptors ^3^. Endogenous eCBs, including anandamide (AEA) ^4^and 2-arachidonoylglycerol (2-AG) ^5,6^, are bioactive lipids derived from polyunsaturated fatty acids. Interestingly, the ECS has been shown to influence macrophage polarization.

Activation of CB2R predominantly promotes anti-inflammatory M2 macrophage polarization ^7-12^, while the roles of CB1R in macrophage polarization remain almost completely underexplored. The underappreciation of CB_1_R’s role in immune regulation highlights the importance of further exploring its involvement. Given the diverse cellular pathways influenced by the ECS, its involvement in the host response to bacterial infections is a topic of interest. Recent review has highlighted the complexity of ECS’s role in this context^13^. Unfortunately, the assessment of CBs’ effects on bacterial infections has been confined to a very limited number of studies^*14-25*^, yielding heterogeneous outcomes contingent on the infection model used. Some infection models show improved outcomes with CB treatment^20^, while others demonstrate immunosuppressive effects, leading to decreased host resistance^19,23,24^. These variable outcomes emphasize the need for further and context-dependent research.

*Salmonella* presents a unique challenge due to its intricate survival strategies within the host. Our previous studies described *Salmonella*’s influence on host lipid metabolism, including eicosanoids^26-28^, which serve as sources of eCBs. We have shown that *Salmonella* infection reduces the activity of α/β-hydrolase domain 6 (ABHD6) and fatty acid amide hydrolase (FAAH), two key enzymes responsible for 2-arachidonoylglycerol (2-AG) degradation in macrophages ^29^. While 2-AG has been shown to enhance the phagocytosis of zymosan particles, its role in *Salmonella* phagocytosis and intracellular survival remains unknown ^29^. Further supporting a potential antimicrobial function, elevated 2-AG levels in a murine model were shown to protect against gastrointestinal infections caused by *Citrobacter rodentium* ^25^, suggesting a direct role for 2-AG in countering bacterial virulence. However, the specific contributions of cannabinoid receptors to host immune responses against *Salmonella* infection remain largely unknown, necessitating further study.

In this study, we examined the roles of CB1R and CB2R signaling in modulating the host response to *Salmonella* infection, aiming to uncover novel, cannabinoid receptor-mediated mechanisms of innate immune regulation. Our findings highlight a role for the endocannabinoid system not only in shaping the gut microbiome but also in orchestrating host innate immune defenses against bacterial pathogens.

## Results

### Analysis of endocannabinoid pathway modulation in Salmonella-infected macrophages

Building on our previous observation that endocannabinoid hydrolases, including FAAH and ABHD6, were downregulated at 18- and 24-hours post-infection (hpi) in macrophages infected with *Salmonella* ^*27*^, we investigated whether the expression of endocannabinoid receptor genes, *cnr1* and *cnr2* (encoding CB1R and CB2R proteins, respectively), followed a similar regulatory pattern. To assess this, we examined Cnr1 and Cnr2 expressions in bone marrow-derived macrophages (BMDMs) isolated from C57BL/6 mice and infected with *Salmonella* at a multiplicity of infection (MOI) of 30. Our results revealed a significant downregulation of Cnr1 as early as 2 hpi, which persisted through 24 hpi (**Fig. S1A**), suggesting rapid suppression of CB1R signaling as part of an early immune response. In contrast, Cnr2 expression remained stable across all time points, indicating a distinct regulatory mechanism for CB2R in *Salmonella*-infected BMDMs under these conditions (**Fig. S1B**). These findings highlight a potential role for CB1R in the host response to *Salmonella* infection, while CB2R may be regulated through post-transcriptional or post-translational mechanisms.

### Cannabinoid receptors affect the recruitment of immune cells during infection

Distinct expression patterns of Cnr1 and Cnr2 during Salmonella infection suggest that CB1R and CB2R play different roles in macrophage responses. To assess their functional impact, we examined immune cell recruitment in vivo using knockout (KO) mouse models. CB1R-KO, CB2R-KO, and wild-type (WT) C57BL/6 mice were orally infected with *Salmonella* (7.5 × 107 CFUs) and analyzed at 4 days post-infection (dpi). Flow cytometry revealed that CB2R-KO mice had a significant reduction in natural killer (NK) cells (**Fig. 1B**), while T cell (CD3^+^) and B cell populations remained unchanged (**Fig. 1C, D**). In contrast, CB1R-KO mice exhibited an increased percentage of T cells and a decreased percentage of B cells (**Fig. 1C, D**). Additionally, CB11b^+^F4/80^+^ macrophages were elevated in CB1R-KO mice but reduced in CB2R-KO mice (**Fig. 1E-G**), whereas neutrophils followed the opposite trend. These findings demonstrate distinct, receptor-specific roles for CB1R and CB2R in immune cell recruitment during *Salmonella* infection.

**Figure 1:**
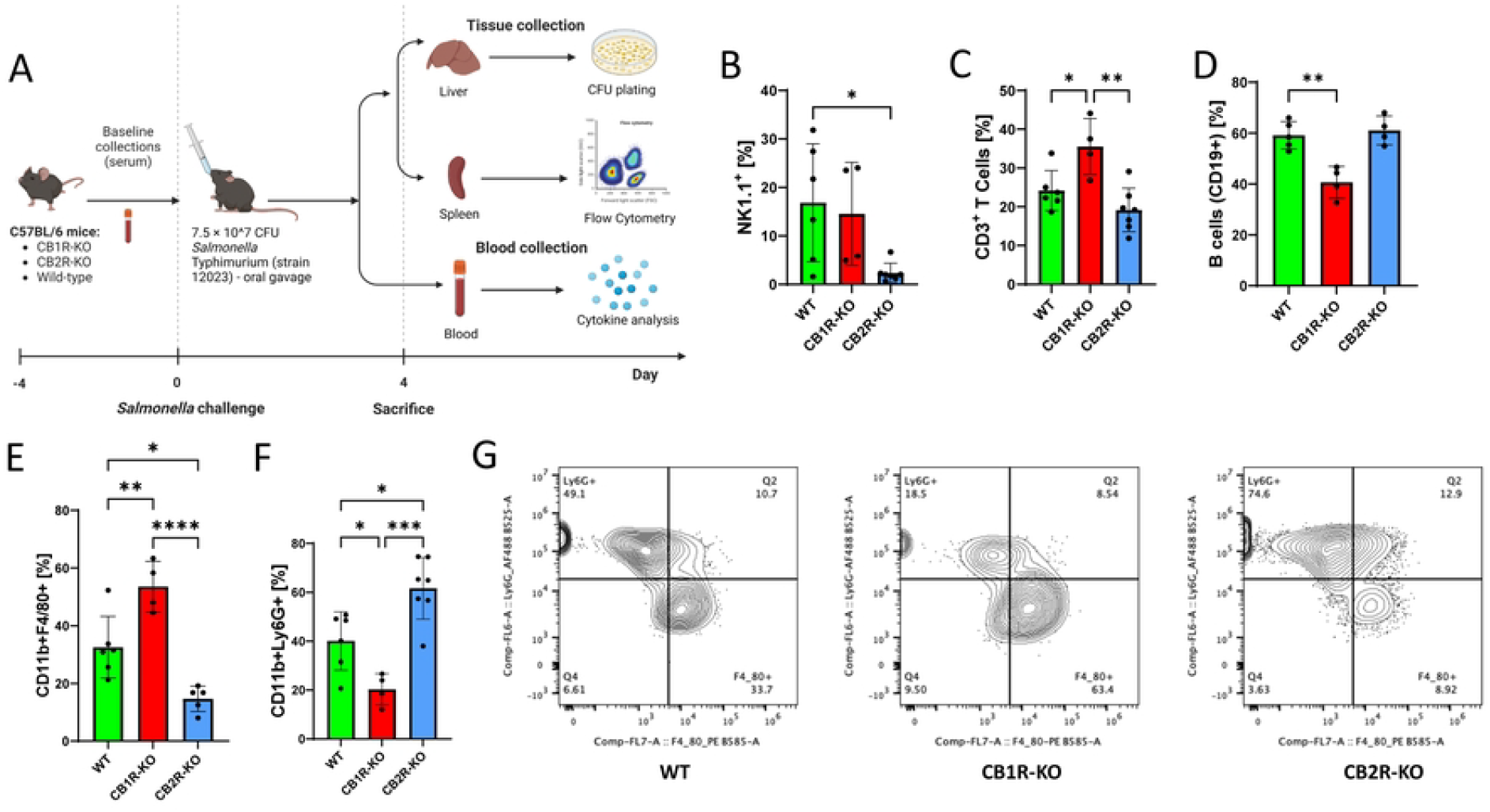
Roles of CB1R and CB2R deficiencies in modulating host resilience and survival against *Salmonella* Typhimurium infection in C57BL/6 mice. **(A)** Experimental setup illustrating the oral infection of wild-type (WT), CB1 receptor knockout (CB1R-KO), and CB2 receptor knockout (CB2R-KO) C57BL/6 mice with 7.5 × 10^7 CFUs of *Salmonella* Typhimurium. **(B–F)** Flow cytometry analysis of various immune cell subsets in the spleens of these mice post-infection, showing variations in the innate immune response across the three genotypes. Quantified cell populations include NK cells **(B)**, CD3+ T cells **(C)**, B cells **(D)**, macrophages **(E)**, and neutrophils **(F)**. One-way ANOVA was used for statistical analysis. Statistical significance is denoted by asterisks: *p < 0.05, **p < 0.01, ***p < 0.001, and ****p < 0.0001. **(G)** Comparison of macrophage and neutrophil populations across WT, CB1R-KO, and CB2R-KO genotypes.

### Macrophage cytokine production and polarization in CB1R and CB2R KO mice

Given the observed differences in immune cell recruitment, we next evaluated cytokine production and macrophage polarization in the spleens of Salmonella-infected mice (**Fig. 2**). Flow cytometry analysis revealed that splenic macrophages from CB1R knockout (KO) mice produced significantly less IL-10 (**Fig. 2A**) while simultaneously expressing higher levels of IL-6 (**Fig. 2B**) and TGF-β (**Fig. 2C**) compared to wild-type controls. In contrast, macrophages from CB2R-KO mice also showed reduced IL-10 production and elevated TGF-β levels (**Fig. 2C**), while their IL-6 levels remained unchanged. Notably, intracellular TNF-α levels were lower in CB1R-KO macrophages and higher in CB2R-KO macrophages (**Fig. 2D**); however, these measurements only reflect intracellular abundance rather than secreted cytokine levels. Furthermore, CB1R-KO macrophages exhibited increased expression of the costimulatory marker CD86 (**Fig. 2E**) without significant changes in CD206 expression. In contrast, CB2R-KO macrophages not only displayed higher CD86 levels **(Fig. 2E)** but also demonstrated a decrease in CD206 expression **(Fig. 2F)**, collectively indicating a shift toward a pro-inflammatory phenotype.

**Figure 2.**
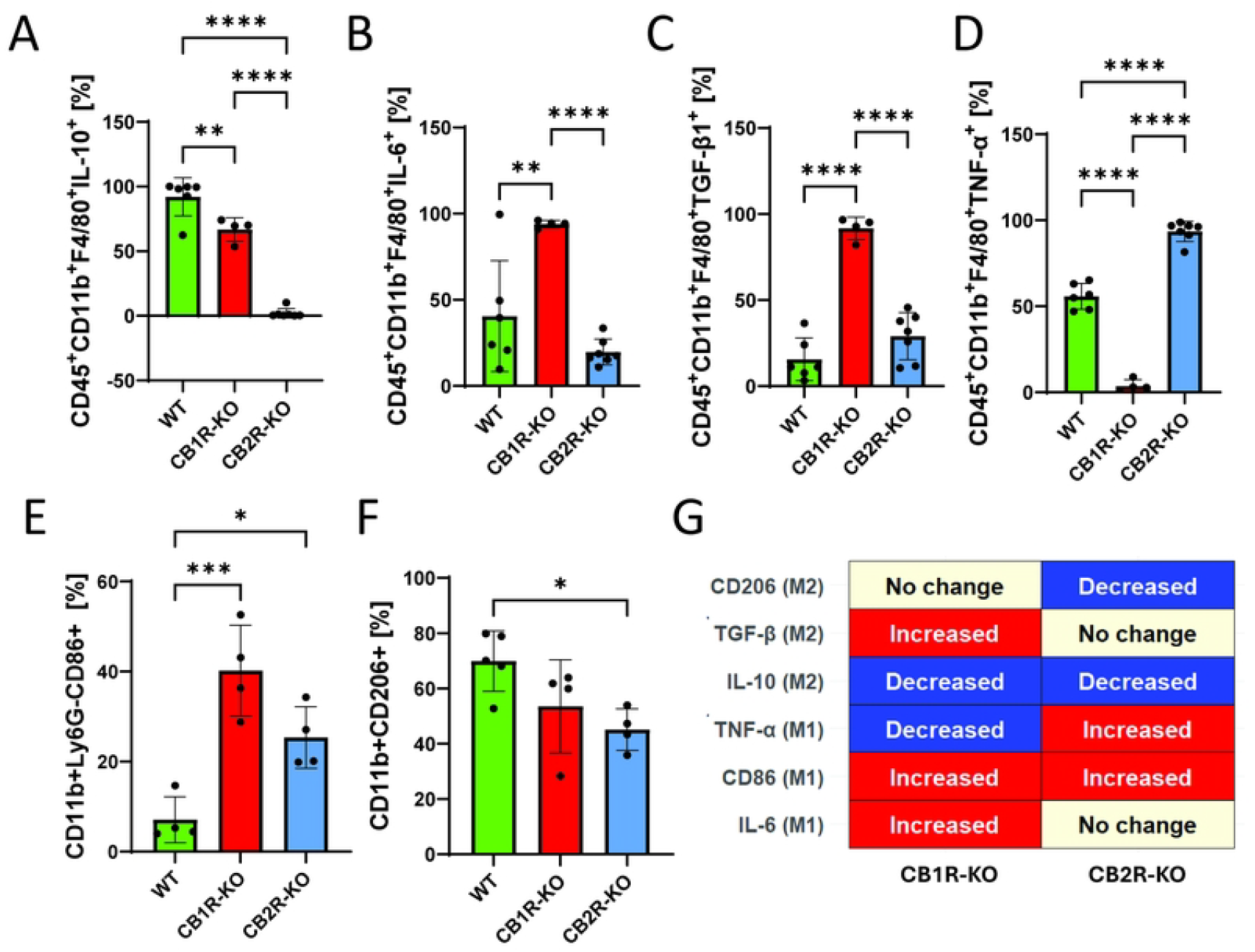
Cytokine expression in splenic macrophages in CB1R and CB2R knockout mice post-infection with *Salmonella* Typhimurium. Flow cytometry was used to quantify cytokine production in splenic macrophages (CD45^+^CD11b^+^F4/80^+^) from wild-type (WT), CB1 receptor knockout (CB1R-KO), and CB2 receptor knockout (CB2R-KO) C57BL/6 mice, 4 days post-infection with *Salmonella* Typhimurium. The following markers were analyzed: **(A)** IL-10, **(B)** IL-6, **(C)** TGF-β, **(D)** CD86, **(E)** TNF-α, and **(F)** CD206. Each bar graph represents the mean percentage ± SEM, with individual data points for each mouse. Statistical analyses were performed using one-way ANOVA to compare cytokine and marker expression levels between WT, CB1R-KO, and CB2R-KO groups. Statistical significance is denoted by asterisks: *p < 0.05, **p < 0.01, ***p < 0.001, and ****p < 0.0001. **(G)** Graphical summary of M1/M2 macrophage polarization markers in CB1R-KO and CB2R-KO splenic macrophages post-*Salmonella* infection, highlighting the receptor-specific shifts in immune phenotypes.

While both CB1R-KO and CB2R-KO macrophages show elements of a pro-inflammatory, M1-like polarization, a closer look reveals distinctions between the two **(Fig. 2G)**. Specifically, CB2R deficiency results in a clear M1 profile with reduced IL-10, increased CD86, and decreased CD206. In contrast, CB1R deficiency leads to a more complex or dysregulated state: despite reduced IL-10 and increased CD86—features consistent with M1 polarization—the concomitant elevation of both IL-6 and TGF-β suggests elements of an M2b-like phenotype. This mixed profile may impair effective bacterial clearance despite the pro-inflammatory context.

Overall, the data suggest that CB1 and CB2 signaling help maintain a balanced, anti-inflammatory (M2) state, and that the loss of either receptor skews macrophages toward a pro-inflammatory phenotype. However, while CB2R deficiency produces a straightforward M1 shift, CB1R deficiency results in a dysregulated polarization that combines aspects of both M1 and M2b profiles.

### Bacterial burden in cannabinoid receptor knockout mice and in vitro

To investigate the role of cannabinoid receptor signaling in host defense against *Salmonella* infection, we assessed bacterial burden in CB1R-KO, CB2R-KO, and WT mice using a sepsis model. Flow cytometry analysis revealed a significant increase in CD45^+^ *Salmonella*^+^ cells in the spleens of CB1R-KO mice (**Fig. 3A, B**) and in all splenocytes (**Fig. 3C**), which correlated with markedly higher bacterial loads in both the liver and spleen of CB1R-KO mice compared to WT controls (**Fig. 3C**). These findings indicate that CB1R deficiency promotes bacterial dissemination, leading to a more severe systemic infection. In contrast, CB2R-KO mice did not exhibit a significant difference in bacterial burden compared to WT controls, suggesting that CB1R plays a dominant role in controlling bacterial proliferation and organ dissemination in this model. These results highlight the critical function of CB1R in host defense against *Salmonella*, likely through its influence on immune cell activation and bacterial clearance mechanisms.

**Figure 3.**
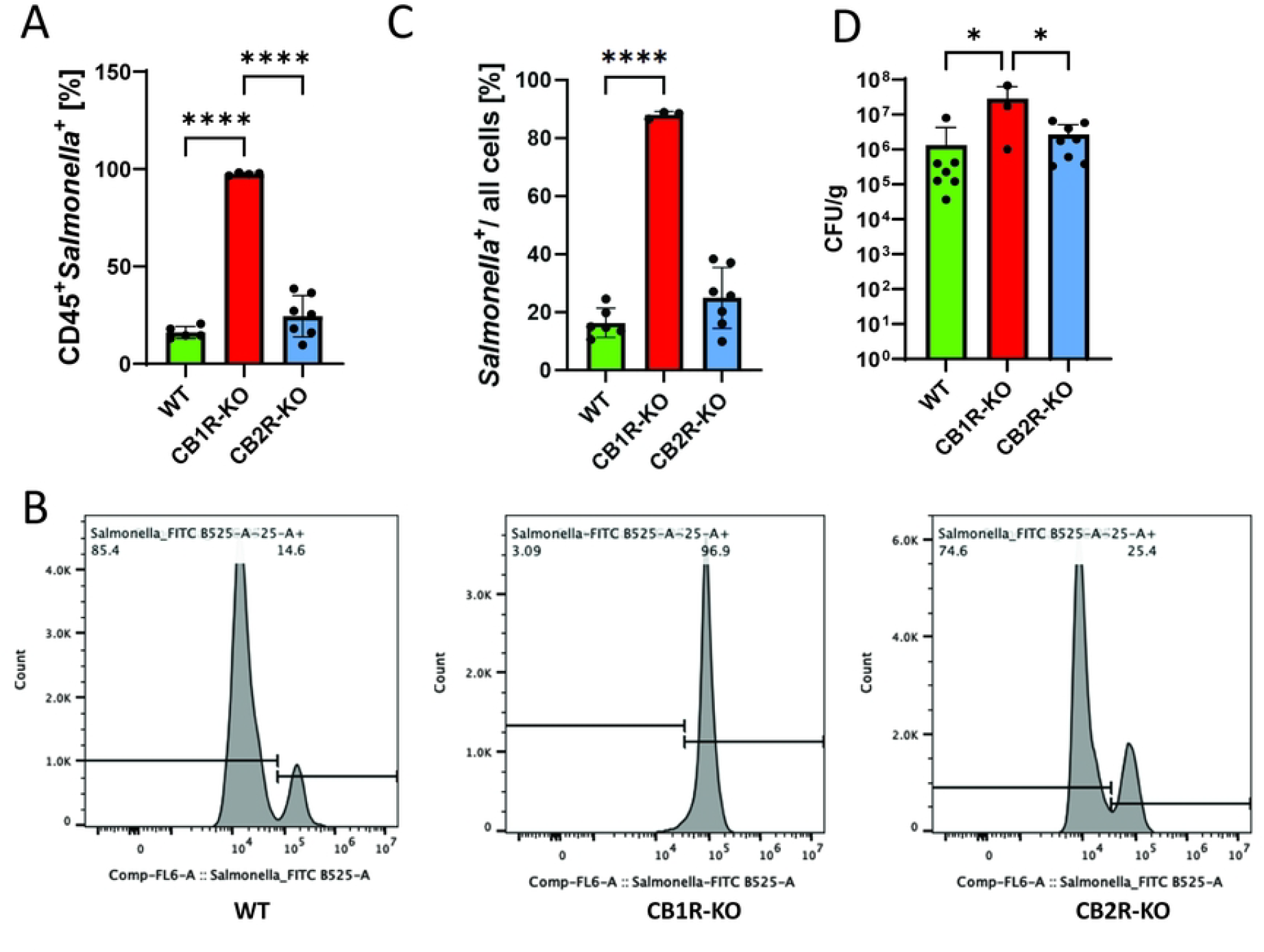
Influence of cannabinoid receptors on *Salmonella* Typhimurium dissemination in C57BL/6 mice. **(A-C)** Flow cytometry analysis of splenic cells from C57BL/6 mice infected with *Salmonella Typhimurium*, distinguishing CD45+ Salmonella+ cells to assess bacterial dissemination. **(A)** presents data from CD45+ cells, while **(B)** expands the analysis to all live cells, mapping the systemic spread of *Salmonella* in CB1R and CB2R knockout (CB1R-KO and CB2R-KO) mice compared to controls. **(D)** Colony-forming unit (CFU) plating of liver and spleen tissues from infected mice to quantify viable *Salmonella* loads. One-way ANOVA was used for statistical analysis. Statistical significance is indicated by asterisks: *p < 0.05, ****p < 0.0001.

Next, to investigate how CB1R and CB2R deficiencies affect bacterial burden at the cellular level, we conducted in vitro infections using BMDMs isolated from CB1R-KO, CB2R-KO, and WT mice. Macrophages were infected with *Salmonella* for 2 or 24 hours, followed by quantification of bacterial presence and total cell counts. DAPI staining was used to determine cell numbers, while GFP fluorescence intensity, normalized to DAPI, was measured to assess bacterial association within macrophages. At 2 hpi, both CB1R-KO and CB2R-KO macrophages exhibited slightly lower bacterial association per cell compared to WT controls (**Fig. S2A**). However, by 24 hpi, this trend reversed in CB1R-KO macrophages, where intracellular bacteria increased compared to WT cells (**Fig. S2B, C**). This finding is consistent with the increased bacterial burden observed in CB1R-KO mice. Cytokine analysis at 6 hpi revealed some change in inflammatory responses.

TNF-α levels were elevated in CB1R-KO macrophages but reduced in CB2R-KO cells, while IL-10 was decreased in both KO groups relative to WT controls (**Fig. S2D, E**).

All these findings suggest that CB1R is a regulator of macrophage-mediated bacterial control and systemic infection severity, pointing towards its putative role in host-pathogen interactions during *Salmonella* infection. A heightened pro-inflammatory phenotype may initially be beneficial for bacterial killing but can ultimately fail to control bacterial growth if not properly regulated, as appears to be the case in CB1R-KO macrophages.

### Host resilience and survival in CB1R and CB2R deficient mice

To assess the impact of CB1R and CB2R deficiencies on host resilience during *Salmonella* infection, we conducted a *Salmonella* challenge study in CB1R-KO, CB2R-KO, and WT mice including outputs such as weight loss monitoring, body condition scoring, serum cytokine analysis, and survival tracking. Groups of mice were orally infected with *Salmonella* Typhimurium (7.5 × 107 CFUs) and monitored for one-week post-infection. Body weight and condition scores were recorded daily, while serum cytokine levels were measured to assess inflammatory responses. Survival rates were tracked for up to four days post-infection to determine overall susceptibility. WT mice exhibited only a moderate decline in body weight and maintained relatively stable body condition scores throughout the infection period (**Fig. 4A, B**). In contrast, both CB1R-KO and CB2R-KO mice suffered significant weight loss and exhibited a notable decline in body condition, indicating increased susceptibility to infection-induced cachexia (**Fig. 4C, D**). Serum cytokine analysis revealed key differences in systemic inflammation. CB1R-KO mice displayed significantly elevated IL-1β and TNF-α levels compared to WT and CB2R-KO mice (**Fig. 4E, F**), suggesting a heightened systemic inflammatory response. This excessive inflammation likely contributes to the increased severity of infection observed in CB1R-KO mice. While CB2R-KO mice also showed elevated TNF-α levels, the increase was less pronounced, suggesting a distinct role for CB1R in modulating inflammatory responses. Survival analysis demonstrated the protective role of CB1R in host defense. WT mice had the highest survival rate (∼80%) over the 4-day post-infection period, whereas both CB1R-KO and CB2R-KO mice showed significantly reduced survival, with CB1R-KO mice exhibiting the most severe vulnerability in this systemic model of *Salmonella* infection (**Fig. 4G**).

**Figure 4.**
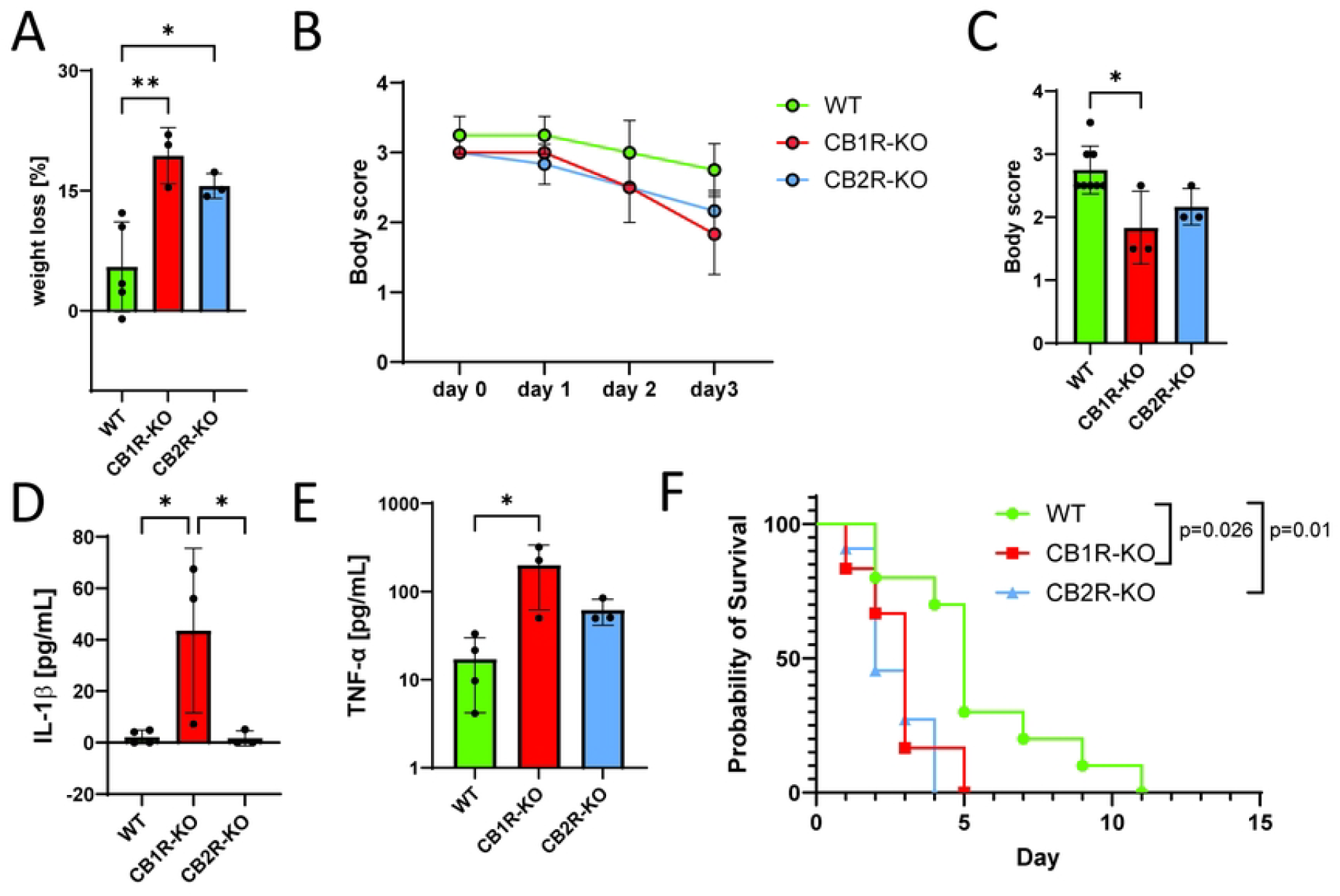
Impact of CB1R and CB2R Knockout on *Salmonella* Infection Outcomes in Mice. **(A)** Body weight loss in wild-type (WT), CB1R knockout (CB1R-KO), and CB2R knockout (CB2R-KO) mice at 4 days post-infection (DPI) with *Salmonella Typhimurium*. **(B)** Body condition scores in WT, CB1R-KO, and CB2R-KO mice at 1, 2, 3, and 4 DPI. **(C)** Body condition scores in WT, CB1R-KO, and CB2R-KO mice at 4 DPI. **(D)** Serum IL-1β levels in WT, CB1R-KO, and CB2R-KO mice at 4 DPI. **(E)** Serum TNF-α levels in WT, CB1R-KO, and CB2R-KO mice at 4 DPI. **(F)** Kaplan-Meier survival curves for WT, CB1R-KO, and CB2R-KO mice over 4 DPI. *Data are presented as mean ± SEM. For panels (A), (C), (D), and (E), one-way ANOVA was used for statistical analysis. Statistical significance is indicated as follows: *p < 0.05, **p < 0.01.

### Host microbiome alterations in CB2R mice during infection

Although our study primarily focused on the role of CB1R and CB2R in immune responses during *Salmonella* infection, we observed that CB2R-KO mice exhibited worse infection outcomes despite showing no significant differences in bacterial dissemination to organs or intracellular bacterial burden in macrophages. This unexpected finding led us to explore potential alternative factors that could contribute to the heightened susceptibility of CB2R-KO mice. Given the well-established role of the gut microbiome in shaping immune responses ^30^ and host defense mechanisms against *Salmonella* ^31-35^, we hypothesized that CB2R deficiency may alter gut microbial composition in a way that may influence susceptibility to infection. The endocannabinoids are known to regulate gut homeostasis and microbiome^36^, making it plausible that loss of CB2R could drive specific microbiome alterations that contribute to immune dysregulation or nutritional landscape. To investigate this possibility, we performed a comparative microbiome analysis in CB2R-KO and WT mice, assessing microbial diversity and composition before and after *Salmonella* infection. The analysis revealed distinct differences in microbiome composition between the two groups, although no significant differences were observed in overall diversity (Shannon diversity index, p=0.32) (**Fig. 5A**). Notably, the abundance of beneficial gut bacteria, including *Lactobacillus acidophilus, L. intestinalis, L. gasseri, L. crispatus*, and *Bifidobacterium animalis*, was significantly reduced in CB2R-KO mice compared to WT controls (**Fig. 5B**). These species are known to promote gut barrier integrity, enhance anti-inflammatory responses, and contribute to pathogen resistance. Conversely, CB2R-KO mice exhibited a significant enrichment of *Alistipes* species (*A. humii, A. finegoldii, A. onderdonkii*), which have been associated with both beneficial and pro-inflammatory effects, metabolic dysfunction, and colorectal cancer (**Fig. 5C**). During infection, pathway enrichment analysis revealed a marked downregulation of key metabolic pathways in CB2R-KO mice, including the *Bifidobacterium* shunt, *L-glutamine biosynthesis III*, and *L-lysine biosynthesis II*, while the *preQ0 biosynthesis* pathway was also significantly decreased compared to wild-type animals (**Fig. 6**). These findings indicate that CB2R deficiency reshapes the gut microbiota during *Salmonella* infection, potentially increasing host susceptibility. Further studies should determine how these microbial shifts impair immune defense against *Salmonella* in CB2R-KO mice.

**Figure 5.**
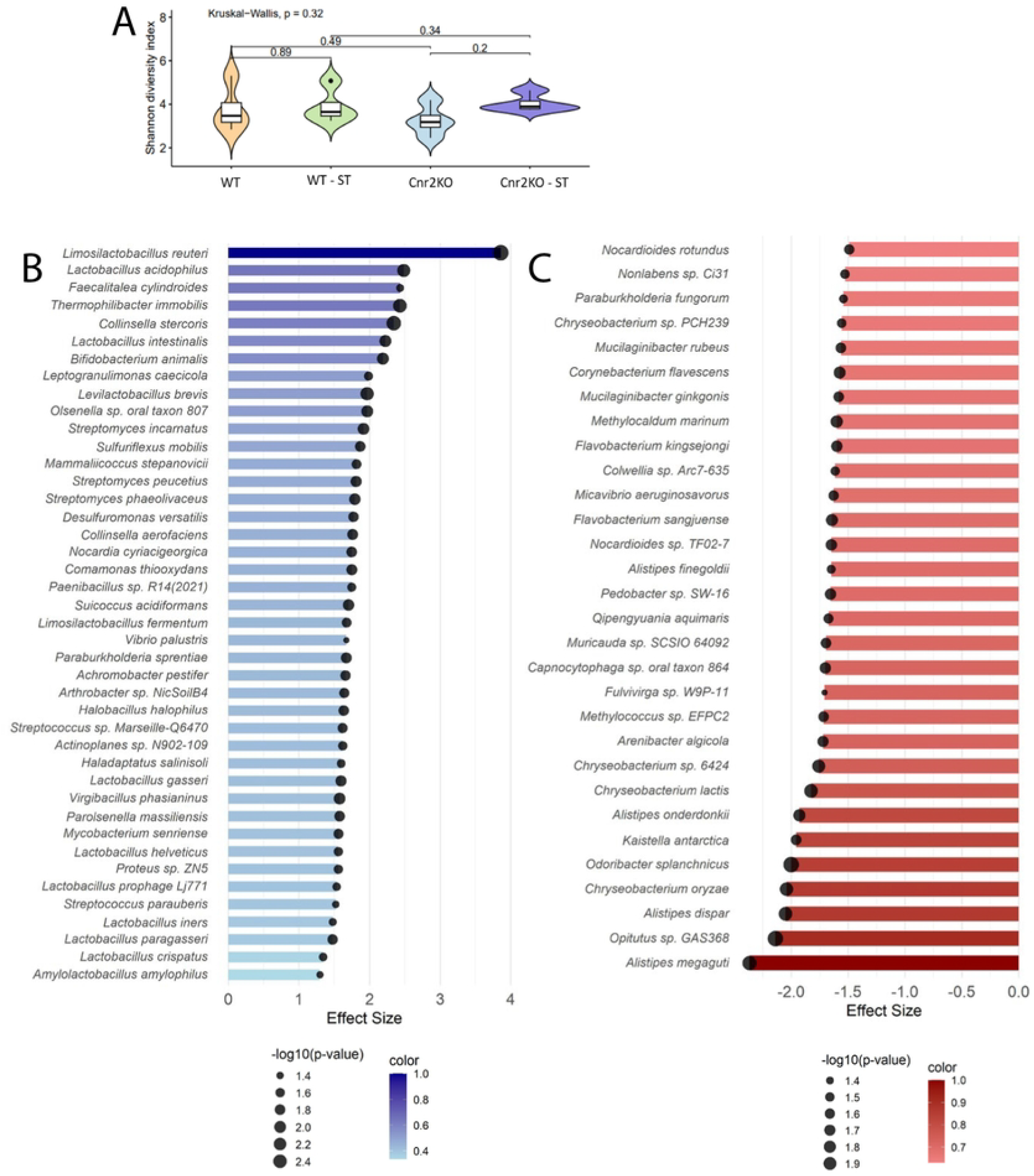
Gut microbiome alterations in CB2R-KO mice during *Salmonella* infection. **(A)** Alpha diversity of the gut microbiota in CB2R knockout (CB2R-KO) and wild-type (WT) mice at baseline (pre-infection) and 4 days post-infection (dpi) with *Salmonella* Typhimurium. Alpha diversity was assessed using the Shannon diversity index. The Kruskal-Wallis test was performed for significance test. **(B-C)** Differential abundance analysis of microbial taxa 4 days post-infection. Blue bars indicate taxa enriched in WT; red bars indicate enrichment in CB2R-KO. Circle size represents -log10(p-value).

**Figure 6.**
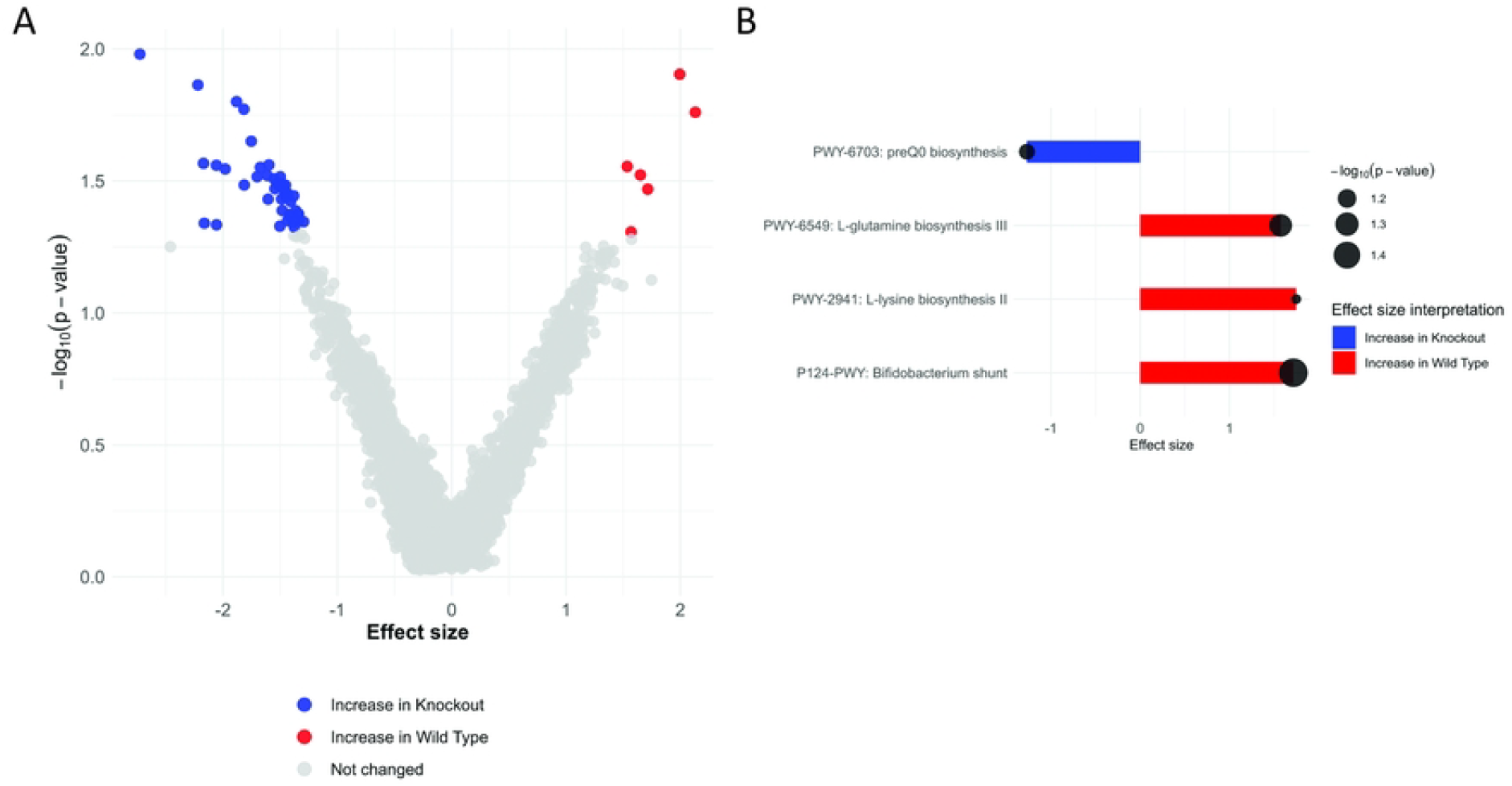
Functional microbiome analysis of CB2R-KO vs. WT mice post-infection. **(A)**. The volcano plot displays differentially abundant metabolic pathways, with red indicating pathways enriched in WT and blue in CB2R-KO. **(B)** Bar plots highlight significantly altered pathways, including downregulation of *Bifidobacterium* shunt activity, L-glutamine, and L-lysine biosynthesis in CB2R-KO mice. Circle size reflects -log10(p-value).

## Discussion

Successful defense against *Salmonella* infection requires a finely tuned immune response— excessive inflammation can lead to host tissue damage, while an insufficient response permits bacterial survival and dissemination ^37^. Our study demonstrates that CB1R and CB2R serve distinct and non-redundant roles in regulating host innate immunity and resilience to *Salmonella* infection. The absence of either receptor results in immune dysregulation and worsened infection outcomes through different mechanisms: CB1R deficiency drives excessive inflammation and weight loss in infected animals, whereas CB2R deficiency disrupts immune homeostasis and induces functional microbiota alterations that may exacerbate infection susceptibility.

### CB1R: a double-edged sword in host defense

CB1R has been extensively characterized in the central nervous system (CNS)^38,39^, but it also modulates immune cells such as macrophages ^40^. Its activation by Δ9-THC, the psychoactive component of *Cannabis sativa* ^41 42^, or endogenous ligands such as AEA and 2-AG ^43^. Mechanistically, CB1R activation suppresses adenylate cyclase activity ^44-46^ and can regulate calcium and potassium channels ^47^, influencing both phagocytic capacity ^40^ and cytokine output ^48^. Although CB1R can promote an inflammatory response under certain conditions ^49,50^, it can also limit excessive inflammation, as evidenced by the effects of agonists like WIN 55212–2 on reducing cytokine production and inflammasome activation ^12 51,52^.

In our study, CB1R deficiency resulted in increased systemic inflammation, pronounced weight loss, and reduced survival in Salmonella-infected mice. Despite this heightened inflammatory backdrop, CB1R-KO macrophages also exhibited a mixed or “M2b-like” phenotype, marked by elevated TGF-β and IL-6 alongside reduced IL-10. This pattern is consistent with M2b macrophages, which can produce both pro-inflammatory (e.g., IL-6, TNF-α) and anti-inflammatory (e.g., IL-10) mediators^53^. However, in our CB1R-KO cells, IL-10 levels were paradoxically lower, suggesting a dysregulated M2b-like state in which certain anti-inflammatory mechanisms (IL-10) are diminished, yet TGF-β is high enough to suppress effective antimicrobial functions (e.g., NO production via iNOS) ^54^. This apparent contradiction—simultaneous hyperinflammation at the systemic level but reduced IL-10 and increased TGF-β at the cellular level—can be explained by the functionally wide-ranging nature of M2b macrophages, which can drive persistent, low-level inflammation while also impairing pathogen clearance ^53,55,56^. Indeed, *Salmonella* has been shown to exploit M2-like phenotypes to establish chronic infections, leveraging changes in lipid metabolism and membrane trafficking ^53^. Our findings thus suggest that CB1R normally helps orchestrate a more balanced macrophage response: in its absence, macrophages become skewed toward a dysfunctional M2b-like state that fails to eradicate bacteria effectively and contributes to overall immune dysregulation. Thus, CB1R signaling plays a dual role in infection control, both promoting bacterial clearance and preventing excessive inflammation. Its absence skews macrophage polarization in a way that favors *Salmonella* persistence, identifying CB1R as a potential target for immunomodulatory therapies.

In addition to modulating macrophage responses, our data suggest that CB1R deficiency may also significantly impair hil recruitment to infected tissues. Although evidence is limited, previous studies have linked CB1R to neutrophil chemotaxis. For instance, activation of CB1R with the agonist ACEA was found to promote neutrophil chemotaxis—a response that can be inhibited by the CB1R antagonist AM281 ^57^. In CB1R-KO mice, the marked reduction of neutrophils in affected tissues indicates that impaired chemotaxis or adhesion may be a key contributor to the observed phenotype. This finding points to an alternative mechanism by which CB1R signaling enhances host defense—not only by modulating macrophage polarization, which could be secondary effect, but also by facilitating effective neutrophil trafficking. Further investigation into the chemokines, adhesion molecules, and signaling pathways driving neutrophil recruitment at specific time points in CB1R-deficient genotype is warranted.

### CB2R deficiency disrupts immune homeostasis and the gut microbiota

CB2R-KO mice exhibited a hyperinflammatory M1a-like macrophage phenotype, marked by increased IL-6 levels and a significant reduction in IL-10. Unlike CB1R-KO mice, CB2R-KO macrophages did not show elevated TGF-β, indicating a lack of compensatory immunosuppression. This reinforces CB2R’s established role in curbing excessive immune activation and promoting an M2 phenotype ^8,9,11,12,20^. Additionally, CB2R-KO mice showed increased neutrophil recruitment, yet this did not enhance bacterial clearance. Instead, excessive neutrophil infiltration likely contributed to tissue damage rather than effective pathogen control, supporting CB2R’s function as a negative regulator of neutrophil-driven inflammation^20^.

Despite no significant differences in bacterial dissemination to organs, CB2R-KO mice exhibited worse infection outcomes, prompting an investigation into alternative contributors to disease severity. Microbiome analysis revealed significant alterations in gut bacterial composition in CB2R-KO mice. Notably, beneficial *Limosilactobacillus reuteri, Lactobacillus acidophilus, L. intestinalis, L. brevis*, and *Bifidobacterium animalis*, which support gut barrier integrity ^58^ and production of anti-inflammatory cytokines ^58 59^, were significantly reduced in CB2R-KO mice after infection. The loss of these protective microbes may compromise gut homeostasis, heightening inflammation and impairing immune tolerance. Conversely, *Alistipes finegoldii, A. onderdonkii*, and *Alistipes megaguti* were significantly enriched in CB2R-KO mice. While *Alistipes* species have been associated with beneficial effects in some diseases, they have also been linked to heightened inflammation, metabolic dysfunction, and colorectal cancer ^60-62^.

Functional microbiome analysis revealed a downregulation of key metabolic pathways, including *Bifidobacterium* shunt activity and amino acid biosynthesis, further supporting the role of CB2R in shaping microbial functional capacity and its potential impact on immune responses. *Bifidobacterium* species and their metabolic products, particularly short-chain fatty acids (SCFAs), are well-documented immunomodulators that enhance gut barrier integrity and pathogen resistance. The Bifidobacterium shunt, a crucial metabolic pathway for SCFA production from oligosaccharide*s* ^63^, is notably depleted in dysbiotic conditions ^64,65^. Similarly, reduced biosynthesis of L-glutamine ^66^, which is essential for epithelial repair, suggests a potential mechanism for impaired mucosal defenses in CB2R-KO mice. Additionally, lysine has been shown to alleviate DSS-induced colitis by reducing inflammation, strengthening the intestinal barrier, and modulating immune cell populations—specifically increasing CD103+ dendritic cells (DCs) and regulatory T cells (Tregs) while decreasing Th1/Th2/Th17 cells ^67^. The observed reduction in L-lysine levels in CB2R-KO mice may contribute to a diminished protective and anti-inflammatory environment, further linking CB2R signaling to immune regulation through microbial-derived metabolites.

Collectively, these findings indicate that CB2R signaling contributes to gut microbiome stability during *Salmonella* infection, and its absence is associated with microbial dysbiosis and altered immune responses. Further investigation is needed to determine whether specific microbial shifts contribute to the heightened disease severity observed in CB2R-KO mice

### Possible therapeutic implications and conclusion

The distinct roles of CB1R and CB2R in regulating immune responses, inflammation, and gut microbiome composition present promising opportunities for therapeutic intervention. Modulating these pathways could potentially provide supporting strategies to improve infection outcomes.

For example, enhancing CB1R activity may help mitigate excessive inflammation, minimize tissue damage, and improve survival during *Salmonella* infection. However, our findings indicate that CB1R deficiency not only exacerbates systemic inflammation but also impairs bacterial clearance by skewing macrophages toward a dysregulated polarization state that combines M1-like features with an M2b-like profile, and by reducing neutrophil recruitment. Thus, therapeutic strategies targeting CB1R must carefully balance immune suppression to avoid creating a permissive environment for pathogen persistence. Notably, prior studies have demonstrated context-dependent effects of CB1R modulation—Δ9-THC, a CB1R agonist, has been shown to influence cytokine production in *Legionella pneumophila* infection by increasing TNF-α and IL-6 levels ^23^ while CB1R blockade with the antagonist SR141716A enhanced macrophage antimicrobial activity in *Brucella suis* infection^21^. Conversely, CB2R activation appears to be more directly linked to restoring immune homeostasis, particularly in settings where excessive inflammation contributes to immunopathology. Our data show that CB2R deficiency leads to gut microbiome dysbiosis and a more straightforward M1-like pro-inflammatory macrophage phenotype, with reduced IL-10, increased CD86, and diminished CD206 expression. These observations suggest that CB2R-targeted therapies could help maintain microbiota-mediated protective mechanisms and mitigate infection-induced dysregulation. Consistent with this view, previous studies have shown that activation of CB2R by compounds such as JWH133 reduces neutrophil recruitment, NF-κB signaling, and NLRP3 inflammasome activation ^20^, supporting its role in preventing excessive immune activation.

## Conclusions

Our study reveals distinct yet complementary roles of CB1R and CB2R in immune regulation and host-pathogen interactions during Salmonella infection. CB1R modulates both the inflammatory response and macrophage polarization, while CB2R plays a key role in maintaining immune homeostasis and gut microbiota balance. The absence of either receptor leads to immune dysregulation, microbial shifts, and worsened infection outcomes. These findings underscore the potential of cannabinoid receptor signaling as a novel factor in host defense and suggest that targeting CB1R and CB2R—potentially in combination with microbiome-based interventions— could offer innovative therapeutic strategies against bacterial infections.

## Materials and Methods

### Bacterial strains

*Salmonella enterica* serovar Typhimurium strain 12023 was used for murine studies and *Salmonella* Typhimurium strain UK-1 was used for cell infections. For murine studies, the bacteria were cultured in Luria-Bertani (LB) Lennox broth at 37°C with shaking until they reached mid-log growth [OD_600_ nm = 0.75]. For cell culture infections, the bacteria were cultured in LB Lennox broth at 37°C with shaking until they reached OD_600_ nm = 0.5. Bacteria were then harvested, washed with phosphate-buffered saline (PBS), and resuspended in PBS to the appropriate concentration for infection.

### Animals

CB1 receptor knockout (CB1R-KO) and CB2 receptor knockout (CB2R-KO) mice on a C57BL/6 background were used for all experiments. Mice were housed under specific pathogen-free conditions with ad libitum access to food and water. All procedures were approved by the Institutional Animal Care and Use Committee (IACUC) at the University of Florida and conducted in accordance with relevant guidelines.

### Salmonella infection in mice

Male and female C57BL/6 mice (8–12 weeks old) from the holding colony were randomly assigned to infection groups. When possible, littermate controls were used to minimize microbiota-related variations in *Salmonella* colonization. Mice were orally infected with *Salmonella* Typhimurium (7.5 × 10^7 CFU) in 50 μL PBS using a gavage needle, while control mice received PBS alone. Infections were monitored for four days post-infection.

Mice were observed daily for clinical signs of infection, including weight loss and body condition, using a standardized scoring system. Weight and body scores were recorded daily for up to 14 days post-infection. To assess survival, mice were monitored for 14 days post-infection. Survival rates were recorded daily. Mice were euthanized if they exhibited severe clinical symptoms, in accordance with IACUC guidelines. Survival curves were generated using the Kaplan-Meier method, and statistical significance was determined using the log-rank (Mantel-Cox) test.

For experiments requiring tissue analysis, mice were euthanized at four days post-infection, and liver and spleen tissues were aseptically collected. Tissues were homogenized in sterile PBS using a TissueLyser LT (Qiagen), and serial dilutions were plated on LB agar. After 24 hours of incubation at 37°C, CFUs were counted to quantify bacterial load in each organ.

### Flow cytometry

Spleens were collected from both infected and control mice, and single-cell suspensions were prepared by mechanically dissociating the tissues through a 70 μm cell strainer. Red blood cells were lysed using RBC Lysis Buffer (Invitrogen), and the remaining cells were incubated with a Live/Dead viability dye (Zombie Aqua) at 4°C for 15 minutes. After washing with FACS buffer (PBS containing 1% BSA), the cells were treated with Fc Block (TruStain fcX anti-mouse CD16/32, BioLegend) for 5 minutes to minimize non-specific binding. Surface marker staining was performed by incubating the cells with an antibody cocktail targeting specific surface markers at 4°C for 30 minutes. Following another wash with FACS buffer, cells were fixed with Cytofix/Cytoperm solution (BD Biosciences) for 15 minutes. Fixed cells were washed twice with Perm/Wash buffer (BD Biosciences) before staining for intracellular markers. Flow cytometric data were acquired using the CytoFLEX flow cytometer (Beckman Coulter) and analyzed using FlowJo software (Tree Star, Inc.).

Flow cytometry was then used to characterize immune cell populations and their functional states in spleens from infected and control mice using multiple staining panels. For the identification of lymphoid and myeloid cell populations, single-cell suspensions were stained with CD45 (Pacific Blue, Cat# 157212) as a leukocyte marker, Zombie Aqua (Cat#77143) for viability, CD3 (PerCP/Cy5.5, Cat# 100217) for T cells, CD19 (APC/Fire750, Cat# 115557) for B cells, CD11b (Alexa Fluor 647, Cat# 101218) for myeloid cells, NK1.1 (Alexa Fluor 700, Cat# 156511) for natural killer (NK) cells, F4/80 (PE, Cat# 123109) for macrophages, and Ly6G (Alexa Fluor 488, Cat# 127625) for neutrophils. A second panel was used to evaluate the activation and polarization of myeloid cells, incorporating CD86 (Pacific Blue, Cat# 105021) as an activation marker for antigen-presenting cells and CD206 (Alexa Fluor 700, Cat# 141733) for M2 macrophages, alongside the same markers for other immune subsets. A separate panel was designed to detect intracellular *Salmonella*, using CD45 and CD11b to gate on leukocytes and myeloid cells, F4/80 to identify macrophages, and a FITC-conjugated *Salmonella*-specific antibody (Invitrogen, Ref # PA1-73020) to detect intracellular bacteria. Cytokine production was assessed using a panel that included IL-6 (PE, Cat# 504504) as pro-inflammatory markers, and IL-10 (Alexa Fluor 488, Cat# 505013) and TGF-β1 (PerCP/Cy5.5, Cat# 141410) as anti-inflammatory markers, in addition to CD45, CD11b, and F4/80 to identify macrophages. All antibodies were purchased from BioLegend, except for the *Salmonella*-specific antibody (Invitrogen).

### Cytokine measurements

Blood was collected from both infected and control mice at the indicated time points via saphenous vein bleeding. Blood samples were centrifugation at 10,000 × g for 10 minutes at 4°C to separate the serum. The samples were then stored in −20C until further use. The levels of TNF-α and IL-1β in the serum were quantified using specific ELISA kits (R&D Biosystems) according to the manufacturer’s instructions.

### BMDM cell culture and infection

Bone marrow-derived macrophages (BMDMs) were generated from mesenchymal stem cells isolated from the hindlimbs of wild-type C57BL/6 mice. Cells were cultured in RPMI 1640 supplemented with macrophage colony-stimulating factor (M-CSF) (25 ng/ml) to promote differentiation into macrophages. The culture medium, including M-CSF, was refreshed every three days, and BMDMs were considered mature on day 7. For infections, BMDMs were seeded at 6 × 105 cells per well in 12-well plates and allowed to adhere for 24 hours before infection.

*Salmonella enterica* serovar Typhimurium strain UK-1 was grown overnight in Lennox LB broth (16 hours, 37°C, shaking). A subculture was established in fresh Lennox LB broth (25 mL) and grown to an optical density (OD600) of 0.5 to ensure bacteria were in the mid-logarithmic growth phase. BMDMs were infected with *Salmonella* at MOI of 10 in incomplete RPMI 1640 for 1 hour. Following infection, cells were washed with PBS and incubated in RPMI supplemented with gentamicin (100 μg/mL) for 1 hour to eliminate extracellular bacteria. The medium was then replaced with RPMI containing gentamicin (25 μg/mL), and cells were incubated until designated time points. Cell pellets were stored in RNAlater for downstream analyses.

### Microbiome analysis

Mice were placed in sterile containers for stool collection, which was performed the day before infection (baseline) and four days post-infection. Stool samples were collected using sterile tools and immediately transferred into sterile Transnetyx tubes containing DNA stabilization buffer.

Samples were temporarily stored at 4°C before being shipped under controlled conditions to Transnetyx for processing the following day. Microbiota profiling was conducted using shallow shotgun whole-genome sequencing on an *Illumina* platform (1×150 bp). Raw sequencing reads were processed to remove low-quality sequences (expected error >0.5) and fragments shorter than 150 bp using Vsearch^68^. High-quality sequences were then taxonomically classified using Kraken 2 ^69^ with the standard database. The resulting contingency table was converted into a phyloseq object for downstream analyses ^70^. To ensure uniform sequencing depth across samples, data were rarefied to a minimum library size of 1,609,000 reads per sample. Alpha diversity metrics were calculated using the microbiome package in R. Boxplots summarizing alpha diversity distributions were generated using *ggplot2* ^*71*^, and statistical significance was assessed using the Kruskal-Wallis test from base R. Differential abundance analysis was performed using the ALDEx2 package ^72^. Sequence files were processed to remove low-quality sequences (expected error >0.5) and sequences smaller than 150bp using Vsearch. The remaining high-quality sequences were classified using Kraken 2 and the standard database. The resulting contingency table was converted into a phyloseq object for downstream analyses. Data were rarefied by the minimum library size of 1,609,000 per sample. Alpha diversity was measured by using the rarefied dataset with the microbiome package. Boxplots summarizing the alpha diversity distribution were plotted by using *ggplot2* R package. The significance of numerical evaluations of alpha diversity was tested with the Kruskal-Wallis from base R. The ALDEx2 package was used to calculate differential abundance. Functional microbiome analysis was conducted with HUMAnN 3.0 to identify shifts in metabolic pathways ^73^.

### qPCR analysis

Cells were collected via cell scraping, resuspended in RNAlater (Thermo Fisher Scientific), and stored at –20°C until further processing. Total RNA was extracted from cell pellets using the RNeasy Mini Kit (Qiagen), following the manufacturer’s instructions. RNA concentration and purity were assessed using a NanoDrop spectrophotometer (Thermo Fisher Scientific).

Complementary DNA (cDNA) was synthesized from 1 μg of total RNA using the iScript Reverse Transcription Supermix for RT-qPCR (Bio-Rad). Quantitative real-time PCR (RT-qPCR) was performed using SsoAdvanced Universal SYBR Green Supermix (Bio-RadX) on a CFX96 Real-Time System (Biorad). Primer sequences for *Cnr1* (Cat #10025636) and *Cnr2* (Cat #10041595) were obtained from the PrimePCR SYBR Green Assay (Bio-Rad). Melt-curve analysis was conducted to verify primer specificity and amplification efficiency. Relative gene expression was calculated using the ΔΔCt method, with β-actin as the internal reference control.

### Statistical analysis

Data are presented as mean ± SEM. Statistical significance between groups was evaluated using Student’s t-test or one-way ANOVA followed by Tukey’s post-hoc test for multiple comparisons, as appropriate. A p-value of <0.05 was considered statistically significant.

## Author Contributions

H.B.: Investigation; Data Curation; Writing – Review & Editing; S.B.: Investigation; Formal Analysis; D.T.: Investigation; Data Curation; Writing – Review & Editing; L.N.L: Formal Analysis; Data Curation; Visualization; L.F.W.R.: Formal Analysis; Data Curation; Visualization; Writing – Original Draft. M.J.F.: Conceptualization; Methodology; Supervision; Writing – Original Draft; Writing – Review & Editing.

## Data Availability Statement

All data generated or analyzed during this study are included in the manuscript and its supporting information files.

## Acknowledgments

M.J.F. was funded by 2021 Research Grants Program of the Consortium for Medical Marijuana Clinical Outcomes Research, which is funded through State of Florida appropriations, as well as R01 AI158749-03 from the U.S. National Institute of Allergy and Infectious Diseases (NIAID). H.A. B. was supported by 5T32AI007110-38 from the National Institute of Allergy and Infectious Diseases.

